# Multi-channel deep learning with intracranial neurostimulation can localize seizure onset zones in humans

**DOI:** 10.1101/2022.02.28.481828

**Authors:** Graham W. Johnson, Leon Y. Cai, Derek J. Doss, Jasmine W. Jiang, Aarushi S. Negi, Saramati Narasimhan, Danika L. Paulo, Hernán F. J. González, Shawniqua Williams Roberson, Sarah K. Bick, Catie E. Chang, Victoria L. Morgan, Mark T. Wallace, Dario J. Englot

**Author notes:** Corresponding author: Graham Johnson, Village at Vanderbilt, 1500 21st Avenue South, Suite 4333, Nashville, Tennessee 37212.

## Abstract

In drug resistant temporal lobe epilepsy, automated tools for seizure onset zone (SOZ) localization using brief interictal recordings would supplement presurgical evaluations and improve care. Thus, we sought to localize SOZs by training a multi-channel convolutional neural network on stereo-EEG (SEEG) cortico-cortical evoked potentials. We performed single pulse electrical stimulation with 10 drug resistant temporal lobe epilepsy patients implanted with SEEG. Using the 500,000 unique post-stimulation SEEG epochs, we trained a multi-channel one-dimensional convolutional neural network to determine whether an SOZ was stimulated. SOZs were classified with a mean leave-one-patient-out testing sensitivity of 78.1% and specificity of 74.6%. To achieve maximum accuracy, the model requires a 0-350 ms post stimulation time period. Post-hoc analysis revealed that the model accurately classified unilateral vs bilateral mesial temporal lobe seizure onset, and neocortical SOZs. This is the first demonstration, to our knowledge, that a deep learning framework can be used to accurately classify SOZs using cortico-cortical evoked potentials. Our findings suggest accurate classification of SOZs relies on a complex temporal evolution of evoked potentials within 350 ms of stimulation. Validation in a larger dataset could provide a practical clinical tool for the presurgical evaluation of drug resistant epilepsy.

## Introduction

Epilepsy affects over 50 million people worldwide, with temporal lobe epilepsy (TLE) being the most common focal epilepsy.^1^ Approximately 30-40% of TLE patients continue to have debilitating seizures despite maximal therapy with anti-seizure medications.^2^ Drug resistant patients may undergo presurgical evaluation ahead of resection, ablation, or neurostimulation therapies. A major goal of presurgical workup is to find the areas of the brain responsible for seizure generation, i.e. the seizure onset zones (SOZs). However, precise localization of SOZs can be challenging with non-invasive modalities such as scalp electroencephalography (EEG), MRI, and PET. Therefore, invasive intracranial monitoring with stereo-EEG (SEEG) is often pursued to provide direct electrographic recordings of seizures to localize SOZs. Monitoring after SEEG implantation often requires long hospital stays of days to weeks to record multiple ictal events.^3^ Thus, it has been proposed that inter-ictal single-pulse electrical stimulation (SPES) of the SEEG contacts to elicit cortico-cortical evoked potentials (CCEP) can help localize SOZs more efficiently.^4–6^

A challenge of interpreting CCEPs using SEEG is that the foundational work in this field was done using subdural electrode grids that measured consistent electrographic phenomena after stimulation, e.g. N1 (10-50 ms) and N2 (50-300 ms) responses.^4^ However, N1 and N2 wave polarity and morphology are defined based on the consistent electrode orientation of subdural electrode grids relative to the cortical surface, and thus orthogonal to cortical pyramidal neurons. In contrast, SEEG has less consistent orientation relative to cortical structures, and translation of N1 and N2 terminology for subcortical gray matter is even more challenging due to heterogenous cytoarchitecture.^7^ Thus, it is difficult to predict the pattern of CCEP wave morphology for any given SEEG contact. Accordingly, most groups rely on coarse metrics for CCEPs in SEEG such as root-mean-squared power.^6,8^ However, these metrics may miss important electrographic features that could help characterize the epileptogenic network. We propose that a multi-channel one-dimensional convolutional neural network (CNN) is well-suited for recognizing variable evoked wave morphology from multiple SEEG contacts simultaneously. This could be a useful tool to delineate whether a given set of CCEPs resulted from an SOZ or non-SOZ being stimulated. Further, by probing various time windows post-stimulation, we can systematically determine which post-stimulation time periods contain the most important classifying features.

## Methods

### Participants and single-pulse electrical stimulation

We collected over 500,000 post-stimulation 900 ms SEEG epochs from 10 patients with drug resistant TLE who underwent presurgical evaluation (**Table 1**). Clinical data were collected through chart review, and seizure outcomes were assigned using the Engel scale.^9^ This study received Institutional Review Board approval and informed subject consents were obtained. We conducted single pulse electrical stimulation (SPES) with every adjacent bipolar pair of contacts in gray matter for each patient. We used a 10 second, 1 Hz, 300 microsecond, biphasic pulse at 3.0 milliamps with a recording sampling rate of 512 Hz.

**Table 1:**
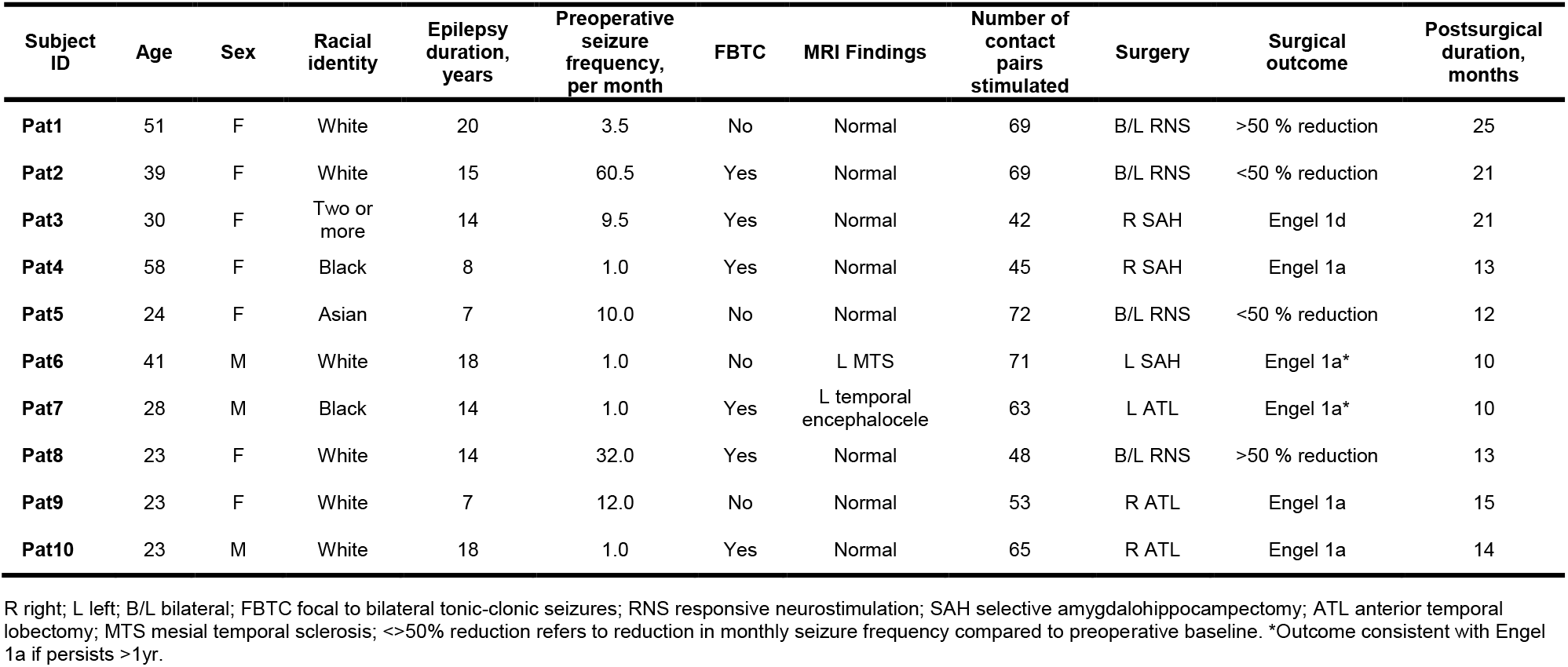
Patient demographics.

### Preprocessing and SOZ labeling

We filtered raw SEEG data using Matlab’s filtfilt function (MathWorks inc., Natick, MA, USA) with Butterworth filter passbands of 1-59, 61-119 and 121-150 Hz. We then parsed the data into 900 ms epochs following each 1 Hz stimulation. This resulted in over 500,000 preprocessed epochs for training our model. SOZs were defined as regions containing any contacts involved in ictal onset of one or more seizures after epileptologist interpretation of all ictal data. Using custom SEEG planning software, CRAnial Vault Explorer (CRAVE), we automatically localized every contact for each patient and created a table of all inter-contact Euclidean distances.^10^

### Deep learning

Using post-stimulation EEG epochs, we trained a one-dimensional multi-channel multi-scale CNN (**Fig. 1A**). To accomplish this, we modified the Multi-Scale-1D-ResNet developed by Fei Wang (https://github.com/geekfeiw/Multi-Scale-1D-ResNet) to input 40 SEEG channels simultaneously. To avoid stimulation artifact and implantation bias, the epochs were distance thresholded to exclude any SEEG channels within 20 mm of the stimulation pair.^11^ For each training pass, we randomized the subset of 40 channels chosen from a patient’s entire available channels. We utilized a weighted binary cross entropy loss function and stopped training after five model epochs. We implemented a leave-one-patient-out testing strategy across all patients. We first tested the ability of the trained model to classify SOZs using the entire 0-900 ms window. Next, we tested the model with only a non-overlapping 50 ms sliding window over the post-stimulation period. We also trained the model on three separate randomized region labels to serve as a control.

**Figure 1:**
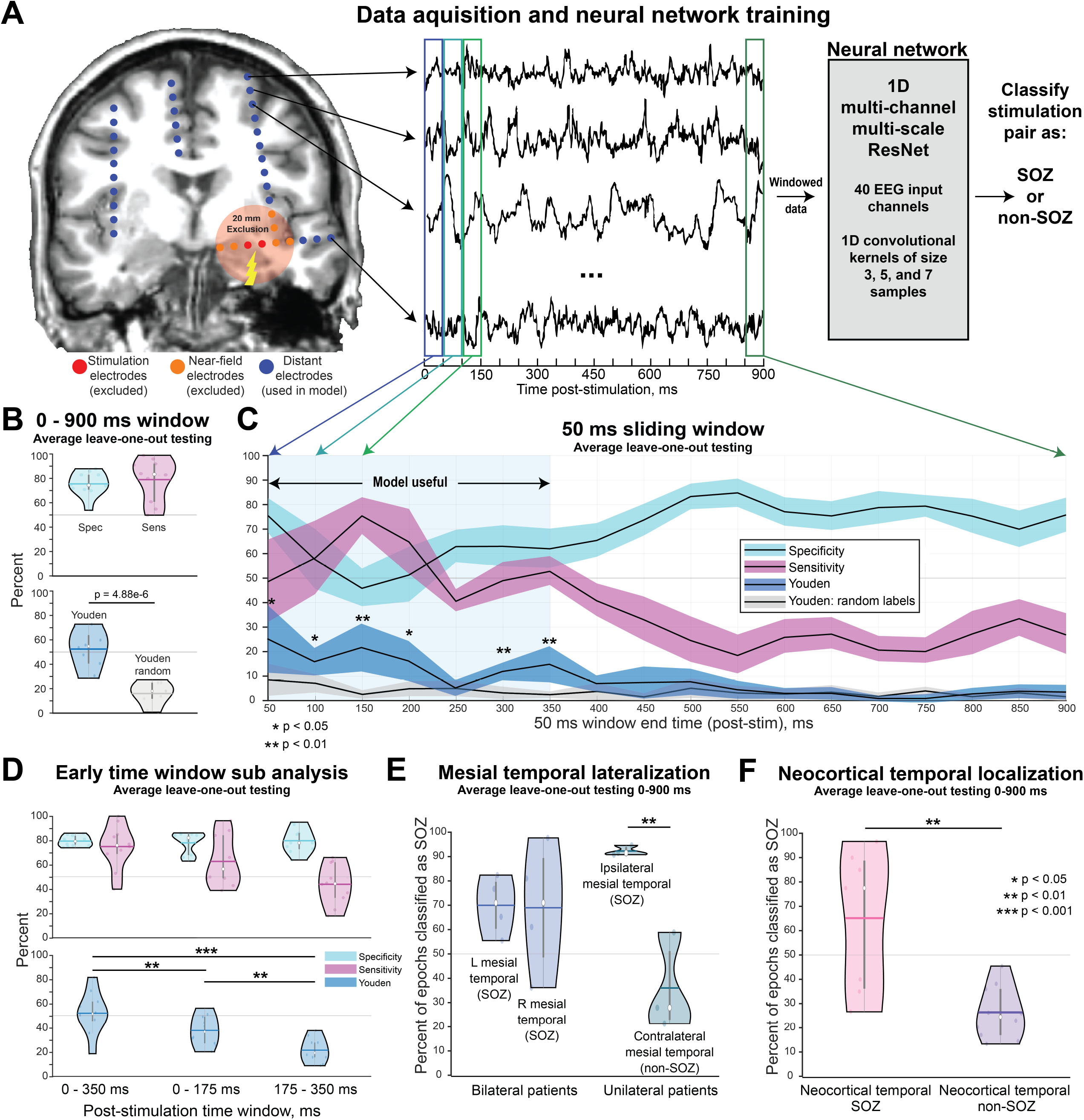
Deep learning on distant SEEG CCEPs can accurately classify SOZs. **A)** We conducted single-pulse electrical stimulation on all gray matter bipolar pairs of contacts for 10 patients undergoing SEEG – this resulted in over 500,000 post-stimulation epochs from the recording channels. To avoid stimulation artifact and biases relating to contact implantation density due to clinical hypotheses, we excluded recordings from contacts that were within 20 mm of the stimulation site. We then trained a convolutional neural network (CNN) to classify if a clinically defined seizure onset zone (SOZ) or non-seizure onset zone (non-SOZ) was stimulated. **B)** We first trained the model using the entire 0-900 ms post-stimulation window. This resulted in a sensitivity of 78.1% and specificity of 74.6% with a significantly improved Youden index compared to training the model on random labels. **C)** For 50 ms sliding windows, the model performed better than random labels for the 0-350 ms post-stimulation 50 ms intervals. Paired t-tests with Bonferroni-Holm multiple comparison correction were conducted between the Youden index (blue), and the random-label Youden index (gray). Note: values on the x-axis represent <u>ending</u> time of the 50 ms window. **D)** Using only the 0-350 ms window resulted in a model accuracy comparable to using the 0-900 ms window. However, dividing this window into 0-175 ms or 175-350 ms resulted in a significant reduction in Youden indexes. **E)** The model was not simply classifying all mesial temporal structures as SOZs. For bilateral patients, the model classified left and right mesial temporal structures as SOZs with comparable confidence around 70%. For unilateral patients, the model correctly classified ipsilateral mesial temporal structures as SOZs at a rate of 91.5% and contralateral (i.e. non-SOZs) at a low rate of 35.1% - suggesting that the model can accurately classify unilateral vs. bilateral seizure onset. **F)** The model was also able to correctly classify neocortical temporal SOZs 64.4% of the time with a low false positive rate of 26.0%. White dot in violin plots is median, horizontal bar is mean.

### Post-hoc testing

We conducted post-hoc testing to determine 1) which post-stimulation time period is best for SOZ classification, 2) can the model classify unilateral vs. bilateral mesial temporal onset, and 3) can the model accurately classify neocortical temporal SOZs? We accomplished (1) by nulling the data outside the desired time window before training. For (2), we calculated the accuracy of left and right mesial temporal SOZ classification for patients with bilateral mesial temporal seizures (n=4) vs. patients with a) unilateral mesial temporal seizures on ictal SEEG, b) a bilateral SEEG implant, and c) seizure-free surgical outcomes (n=3). To accomplish (3), we calculated the accuracy of neocortical temporal SOZ classification in all patients.

### Statistical methods

We calculated the sensitivity and specificity for the leave-one-patient-out testing across all 10 patients for the various time window analyses. We also report the Youden index (sensitivity + specificity – 100) to summarize the usefulness of the model at a given time window; Youden index values above 50 are generally considered to be a very useful model for classification, and values close to 0 are considered useless even if sensitivity or specificity is individually high.^12^ We compared Youden indexes with paired t-tests using Bonferroni-Holm multiple comparison correction.

## Data availability

Data and computer code are available upon reasonable request.

## Results

### CNN trained on long-range CCEPs accurately classifies SOZs

As outlined in **Fig. 1B**, the CNN trained on the entire 900 ms post-stimulation period correctly classified the stimulation pair as SOZ with a mean leave-one-patient-out testing sensitivity of 78.1% (95% confidence interval [CI] 67.8 to 88.4%) and specificity of 74.6% (95%CI 68.7 to 80.5%), resulting in an average Youden index of 52.7 (95%CI 43.7 to 61.8). In comparison, when the model was trained using regions randomly labeled as SOZ or non-SOZ, the average Youden index was significantly decreased to 16.5 (95%CI 9.62 to 23.4, t-test p=4.88e-6). Furthermore, the model achieved significantly improved Youden indexes when training on 50 ms sliding windows ranging from 0-350 ms compared to the same windows with random labels (**Fig. 1C**). Interestingly, the specificity and sensitivity of the model peaked during different periods in the initial 350 ms post-stimulation – the sensitivity peaked for the time window that spans 100 to 150 ms, while the specificity peaked for the 0-50 ms time window. This suggests that delayed responses are most *sensitive* for classifying SOZs, whereas early responses are most *specific* for classifying SOZs.

### Important features are temporally distributed within the initial post-stimulation window

We performed post-hoc analyses to assess which early post-stimulation window was most effective at classifying if an SOZ was stimulated (**Fig. 1D**). Using a time window of 0-350 ms we observed a leave-one-patient-out average testing sensitivity of 74.0% (95%CI 63.3 to 84.7%) and specificity of 78.5% (95%CI 75.9 to 81.1%) with an average Youden index of 52.5 (95%CI 42.1 to 62.9) – very similar to when the model was trained on the entire 0-900 ms. When we divided the 0-350 ms period into 0-175 ms and 175-350 ms, the leave-one-patient-out testing Youden index dropped significantly, suggesting both early and late portions of this time window contribute to model performance.

### The model can classify unilateral vs. bilateral onset mesial temporal lobe epilepsy, and can detect neocortical temporal SOZs

We observed that the bilateral onset patients had left mesial temporal structures correctly classified as SOZs for 68.9% (95% CI 58.7 to 79.1%) of the CCEP epochs, and right mesial temporal structure epochs classified as SOZs for 67.9% (95%CI 45.4 to 90.4%) (**Fig. 1E**). For unilateral patients, the model correctly classified mesial temporal structures ipsilateral to the seizure onset hemisphere as SOZs for 91.5% (95%CI 89.7 to 93.3%) of the epochs with a low false positive rate of 35.1% (95%CI 16.7 to 53.5%) for non-SOZs on the contralateral side. This sub analysis provides evidence that the model was not simply classifying all mesial temporal structures as SOZs, but rather provides accurate classification for unilateral vs. bilateral mesial temporal onset patients. Furthermore, the model correctly classified neocortical temporal SOZs at a rate of 64.4% (95%CI 44.3 to 84.5%), and misclassified neocortical temporal non-SOZs at only 26.0% (95%CI 19.7 to 32.3%) (**Fig. 1F**).

## Discussion

We have demonstrated that a CNN trained entirely on SEEG-derived CCEPs farther than 20 mm from the site of stimulation can classify an SOZ with high sensitivity and specificity in TLE. A strength of this approach is that the model accurately classified SOZs despite the variable morphology of CCEPs during stimulation of SEEG electrodes. Further, the most important post-stimulation features for classification are contained within 0-350 ms. This is not surprising considering that most previous findings using RMS have centered around N1 and N2 responses within 300 ms.^4,13,14^ However, separating this window into smaller segments significantly reduces model accuracy. This suggests that there is a complex pattern of CCEPs occurring at various periods post-stimulation that must be considered in an ensemble to accurately classify the stimulation of ictogenic tissue – this observation could be due to varied phenotypes of evoked responses.^15^ Additionally, through our sub analyses, we conclude that this model was not classifying all mesial temporal structures as SOZs and can accurately distinguish unilateral vs bilateral mesial temporal onset. Finally, the model can also accurately classify neocortical temporal SOZs.

### Limitations and future work

Although 500,000 non-overlapping SEEG epochs were used to train the CNN, training and testing datasets were divided at the patient level. Thus, our relatively small sample size of 10 patients limits our assessment of generalizability and motivated our conservative strategy of leave-one-patient-out testing across the entire cohort. Also, mean follow-up was 15.4 months, and future seizure recurrences may decrease the confidence in clinical SOZ localization and change labels for the CNN. We also did not include any focal extratemporal-lobe epilepsy patients and thus cannot comment on the extension of these techniques to that population. Our future work is aimed at addressing these limitations by collaborating with other institutions that collect these rare datasets. We also hope to test this model on patients with surgical outcomes of Engel 2-4. Perhaps, previously unidentified SOZs, including bilateral seizure onset, could be elucidated in Engel 2-4 patients with a model trained on Engel 1 patients.

### Conclusions

This work serves as the first demonstration, to our knowledge, that a one-dimensional multi-channel multi-scale CNN can learn highly non-linear features of SEEG-derived CCEPs occurring across multiple SEEG channels simultaneously to classify when an SOZ is stimulated. Furthermore, we demonstrated the importance of utilizing the entire 0-350 ms time window for classification. We hope that future work will consider using deep learning as a tool to explore the complex CCEPs generated with SEEG.

## Acknowledgements

We would like to thank the patients participating in our investigation.

## Author contributions

G.W.J. contributed to project conception, data collection, data preprocessing, data analysis, and manuscript preparation. L.Y.C. contributed to data analysis and manuscript preparation. D.J.D. contributed to data analysis, and manuscript preparation. J.W.J. contributed to data collection, data preprocessing, data analysis and manuscript preparation. A.S.N. contributed to data preprocessing, data analysis, and manuscript preparation. S.N., D.P., H.F.J.G., S.W.R., S.K.B., C.E.C., V.L.M., M.T.W. contributed to data analysis and manuscript preparation. D.J.E. contributed to project conception, data analysis, and manuscript preparation.

## Ethics Statement

We confirm that we have read the journal’s position on issues involved in ethical publication and affirm that this report is consistent with those guidelines.

